# Cholinergic Modulation of Neurovascular Coupling and Neuroimaging Signals

**DOI:** 10.1101/2023.11.08.566348

**Authors:** Gaia Brezzo, Jason Berwick, Chris J Martin

## Abstract

The use of non-invasive functional neuroimaging methods to discriminate levels or patterns of brain activity between different groups, across different time points, or in response to drug treatment, depends on an understanding of neurovascular coupling relationships and how they themselves are affected by the factors of interest. For instance, Alzheimer’s disease (AD) features cerebral blood flow (CBF) and neurovascular unit changes as well as being characterised by a cholinergic dysfunction which impacts upon CBF regulation. The present study investigated the acute neurovascular effects of cholinergic agonist and antagonist treatment in a rat model utilising a multimodal measurement approach. Drugs were administered to rats during concurrent imaging of cortical CBF and recording of local field potential responses to somatosensory stimulation. Analysis of concurrent neuronal and vascular measures revealed a pronounced loss of neurovascular coupling following treatment with the cholinergic antagonist scopolamine. Separate analysis of CBF and neuronal responses reveal an interaction effect of stimulus input and drug treatment for both cholinergic agonist and antagonist treatment, as well as an augmentation of both neuronal and haemodynamic response magnitudes after treatment with the clinically prescribed cholinergic agonist donepezil. These findings have implications for the use and interpretation of functional neuroimaging data acquired in individuals with disease-related or pharmacological manipulations of cholinergic function.

## Introduction

The brain’s dependence on its blood supply to adequately provide glucose and oxygen for its functioning has been extensively documented in the literature. Consequently, a deficiency in blood supply leads to decreased or impaired functioning. Brain tissue has a high energetic demand and the brain’s inability to store energy means that a tightly regulated mechanism for continuous adjustment of local cerebral blood flow (CBF) based on neuronal activation is required: this mechanism is termed neurovascular coupling. This system is responsible for the delivery of oxygen, glucose, elimination of toxic substances and for maintaining brain homeostasis (*1, 2*), whilst providing the physiological basis for non-invasive neuroimaging signals as utilised in blood oxygen level dependent (BOLD) functional magnetic resonance imaging (fMRI).

In recent years is has been shown that a number of brain diseases, including Alzheimer’s disease (AD), are associated with pathological changes in blood flow regulation and the neurovascular apparatus that supports it causing neurovascular uncoupling (*3, 4*). Human neuroimaging studies have shown that reductions in CBF correlate with cognitive impairment (*5*), as well as being a risk factor in the development of AD later in life (*6, 7*). An fMRI study (*8*) has shown that task associated increases in CBF are delayed in mild cognitive impairment (MCI) patients and that these deficits become more prominent in AD patients, whilst multimodal MRI techniques have recently revealed impairment of neurovascular unit function in early AD (*9*). These findings implicate altered CBF regulation as an early event in the development of brain pathology and research is thus beginning to investigate how interventions to improve neurovascular function could provide a degree of neuroprotection (*10*).

An important neuropathological component of AD is loss of cholinergic innervation in the basal forebrain and medial septum as well as loss of projection fibres to the neocortex and the hippocampus from the nucleus basalis and medial septum respectively (*11*). Given the known vascular and neurovascular changes observed in AD, an important question is how disease related or therapeutic alterations in cholinergic function affect CBF regulation and neurovascular coupling (*12*), as well as the functional neuroimaging signals which depend on these neurobiological processes. Disruption of the cholinergic system may lead to alterations in neurovascular coupling because of the known vasoactive properties of ACh (*5, 11*). Similarly, pharmacological treatment of AD symptoms using cholinergic drugs may alter neurovascular coupling, impacting upon both preclinical and human clinical fMRI or positron emission tomography (PET) research studies that use haemodynamics as proxy imaging measures of neuronal activity. This question has been partly addressed by two recent studies carried out in rodent models. Firstly, a preclinical pharmacological MRI (phMRI) study (*13*) has suggested that cognition enhancing effects of cholinergic (agonist) drugs, as well as the potential haemodynamic (imaging) biomarkers for these effects, were mediated by their vascular and not neuronal actions, as a peripherally acting cholinesterase inhibitor (neostigmine) was shown to reverse both the fMRI BOLD signal and the memory disturbing effects of the muscarinic antagonist scopolamine in rats despite its inability to cross the blood brain barrier. A second study (*14*) investigated the neurovascular effects of cholinergic manipulations, including measurements of neuronal and haemodynamic responses to sensory stimulation. Here, it was shown that ACh tone modifies both the magnitude of neuronal and hemodynamic responses to sensory stimulation as well as the strength of the correlation between these measures.

Several important questions remain. Firstly, the haemodynamic and neurovascular effects of the most widely prescribed treatment for AD, the cholinergic agonist donepezil, have not been evaluated in this context. Secondly, the assessment of neurovascular function by Lecrux et al. (2017) relies on long-duration stimuli to elicit a ‘steady-state’ neuronal/vascular response, whereas in assessing dynamic neurovascular coupling it is advantageous to utilise brief stimuli so as to inform on the initial neurovascular impulse-response function independent of longer-latency non-linearities in coupling (*15, 16*). This is especially important for the analysis and interpretation of BOLD fMRI studies where the short-latency impulse response function serves as the basis for the canonical model of the hemodynamic response (*17*). Thirdly, because it has previously been shown that anaesthesia used in preclinical studies can affect stimulus-response and neurovascular coupling parameters (*18, 19*), it is important to establish the neurovascular coupling relationship using a wider range of stimulation ‘inputs’, rather than a unitary stimulus type so as to provide improved assurance that findings are robust beyond specific stimulation parameters and experimental (anaesthetic) conditions (*20*).

The present study was designed to identify the effects of the widely prescribed cholinesterase inhibitor donepezil upon haemodynamic responses and neurovascular coupling, with comparison to the effects of the cholinergic antagonist scopolamine. To investigate this we concurrently imaged CBF changes and recorded neuronal activity in the rat somatosensory cortex whilst manipulating functional activation using a range of sensory stimuli.

## Materials and Methods

The present study was approved by the UK Home Office under the Animals (Scientific Procedures) Act 1986 and the University of Sheffield Animal Welfare and Ethical Review Body (AWERB, local ethics committee). All procedures were conducted under a U.K. Home Office licence and have been reported in accordance with the ARRIVE guidelines.

### Animals and pharmacological treatment

Female Hooded Lister rats (*n* = 18, 220g-320g, 4-5 months of age) kept in a 12-hour light/dark cycle environment at a temperature of 22 ⁰C with access to food and water *ad libitum* were housed in polycarbonate cages (*n*=3 per cage) under pathogen free barrier conditions in the Biological Services Unit at the University of Sheffield. Animals were fed conventional laboratory rat food. The animals were randomly assigned to one of three groups (control, scopolamine or donepezil, *n* = 6 per group). Haemodynamic and electrophysiological data were concurrently acquired in all treatment groups in a repeated measures design. Each animal received an intravenous injection based on drug condition: scopolamine animals received 2mg of scopolamine (Tocris, UK) prepared in 1ml of saline; the donepezil animals received a dose of 2mg/kg of donepezil (Tocris, UK) dissolved in saline; control animals were administered a saline vehicle (1ml).

### Surgical procedures

Details of surgical and experimental paradigms were similar to those reported in previous publications (*21-24*). Rats were anaesthetised with an intraperitoneal injection (i.p.) of urethane (1.25mg/kg in 25% solution), with additional doses of anaesthetic (0.1ml) administered if necessary. Choice of anaesthetic was determined by urethane’s suitability for invasive surgery as well as long-lasting stability which makes it well suited to experiments where data collection lasts several hours and where stable neuronal activity (*25*) is required. Moreover, there is experimental evidence that urethane has minimal effects on acetylcholine release (*26*). Route of injection was selected as i.p injection which achieves a deep state of anaesthesia in a short duration (*27*). Anaesthetic depth was determined by means of hindpaw pinch-reflex testing. A topical anaesthetic (Xylocaine, 10mg Spray, AstraZeneca Ltd., UK) was additionally administered if necessary during surgical procedures. Animals were tracheotomised to allow artificial ventilation and regulation of respiratory parameters. The left femoral artery was cannulated to enable mean arterial blood pressure (MABP) monitoring and blood gas sampling/analysis. The left femoral vein was cannulated to allow continuous administration of phenylephrine in order to maintain blood pressure within a physiological range (100-120mmHg). The right femoral vein was cannulated to allow drug administration.

Animals were placed in a stereotaxic frame (World Precision Instruments Inc., USA) for the remainder of the surgical procedure and data collection. To enable CBF recording, the skull was exposed via a midline incision and a section overlaying the left somatosensory cortex (barrel cortex) was thinned to translucency with a dental drill. This section was located 1-4mm posterior and 4-8mm lateral to Bregma (*28*). A thinned skull was typically between 100 and 200 µm thick with the surface vasculature clearly observable. Care was taken during thinning to ensure that the skull remained cool by frequently bathing the area with saline. At the end of the experiments animals were euthanised with an overdose of pentobarbital and cervical dislocation.

### Physiological measurements

Temperature was maintained at 37 ⁰C (± 0.5 ⁰C) throughout surgical and experimental procedures with the use of a homoeothermic blanket and rectal temperature probe (Harvard Apparatus, USA). Animals were artificially ventilated with room air using a small animal ventilator (SAR 830, CWE Inc, USA): the breathing rate of each animal was assessed and modified according to each individual animal’s blood gas measurements. Respiration rates of the animals ranged from 58 to 70 breaths per minute. Blood pressure was monitored during the experiment with a pressure transducer attached to a 5ml plastic syringe containing heparinised saline (Wockhardt, UK, 50 units of heparin per ml). To ensure normoxia and normocapnia, arterial blood from the femoral artery was allowed to flow back from the cannula into a cartridge (iSTAT CG4+, Abbott Point of Care Inc., USA) and blood gases were analysed using a blood gas analyser (VetScan, iSTAT-1, Abaxis, USA). Physiological parameters were within normal ranges throughout the experiment (mean values: PO₂ =78mmHg (±6.66); PCO₂ =36mmHg (±2.41); SO₂ =96%(±1.07)). Total volume of arterial blood extracted at one time did not surpass 95µL. Phenylephrine (Sigma, Aldrich, 0.2-0.8mg/hr) was administered into the left femoral vein using a syringe pump (Sp200i, World Precision Instruments Inc., USA) to counteract reduced blood pressure caused by anaesthesia; blood pressure was maintained between 100-120mmHg.

### Laser speckle contrast imaging

CBF data were acquired through a laser speckle contrast imaging camera (Full field Laser Perfusion Imager (FLPI-2), Moor Instruments, UK) which was positioned above the thinned cranial window. Images were acquired at 25Hz at a spatial resolution of approximately 10µm/pixel. Total recording time varied depending on stimulation paradigm but a 60 second baseline data acquisition period was acquired in each paradigm to obtain a measure of baseline blood flow.

### Electrophysiology

Local field potentials (LFPs) were measured using a multichannel-channel electrode (NeuroNexus Technologies) inserted through a small burr hole made in the previously thinned cranial window. The electrode was inserted perpendicular to the cortical surface under stereotaxic control. Data were acquired at 10 kHz using a preamplifier unit that was optically coupled to a data acquisition device (Medusa, Tucker-Davis Technologies). Experimental control was implemented using custom-written Spike2 code running on a PC attached to a Micro1401 controller unit (Cambridge Electronic Design, UK). This software also controlled stimulus delivery which was directly synchronised to the laser speckle camera.

### Experimental design

One hour after the completion of surgical procedures and once physiological parameters (blood pressure, blood gases) were within the normal range, data acquisition commenced. Experimental runs began with a 60s acquisition of baseline CBF and electrophysiological data, followed by repeated stimulation trials (see below). Data were acquired prior to (‘*pr*e’) and after (‘*post’)* administration of cholinergic drugs (or saline control) 90 minutes after the start of the experiment. Data were acquired within a ferromagnetic cage and upon an air suspension (isolated) workstation (Vision Isostation, Newport Corporation, USA).

### Stimulation paradigms

Stimulation of the whisker pad was delivered via two subdermal stainless steel needle electrodes (12mmx0.3mm, Natus Neurology Incorporated, USA) directly inserted into the whisker pad in a caudal direction which transmitted an electrical current (1.0 mA). For multifrequency stimulation, the whiskers were stimulated at each of six frequencies (1, 2, 5, 10, 20 & 40Hz), for 2s with a stimulus pulse width of 0.3ms. The order of stimulation frequencies was pseudorandomised with 10 trials at each frequency and an inter-trial interval (ISI) of 25s. For long duration stimulation a 16s stimulus (at 10Hz, 10 trials, 60s ISI) was used. The electrical current is generated by an independent amplifier (Isolated Stimulator DS3, Digitimer Ltd., UK) which directly attaches to the electrodes. This intensity has been shown to evoke a robust haemodynamic response without altering physiological factors such as blood pressure and heart rate.

### Data processing and analysis

All (*n* = 18) animals’ data were processed and included in final statistical analysis. Data were processed in Matlab (2016a) using custom written code. Imaging data were spatially smoothed and then analysed using SPM (*29*), with regions of interest (ROIs) selected from a thresholded activation map and expected to include contributions from arterial, venous and parenchymal (capillary bed) compartments. ROI size was consistent across animals, with no significant difference between groups (*F*(2,17,) = 1.926, *p*=.180). Time series of changes for each stimulation trial were then extracted from the ROI and down-sampled to 5Hz (from 25Hz). Data from each stimulation trial were extracted and divided by the average perfusion value (arbitrary units) over a pre-stimulus baseline period (10s), to yield a measure of percentage change in CBF. Time series were averaged across trials according to stimulation condition. The maxima and area under the curve (AUC) for each response were found.

Neuronal recordings for computing local field potential responses were similarly extracted from a single channel of the multichannel electrode that corresponded to a cortical depth of ∼500µm (layer IV), and averaged across trials. AUC measurements were calculated from the response to each pulse in the stimulation train (0-50ms period, with respect to the stimulus pulse) and summed to quantify neuronal response magnitudes. EEG analysis of neuronal data was conducted by computing the power spectra for epochs of electrophysiological data extracted from the multichannel electrode (layers 1 to 4, depth 100-800µm), with power summed cross the channels in five frequency bands corresponding to the EEG delta (0.5-4Hz), theta (4-8Hz), alpha (8-13Hz), beta (13-30Hz) and gamma (30-70Hz) range.

## Results

Materials, data and associated protocols are available on request.

### Baseline blood flow and neuronal activity

To assess any effects of drug administration upon baseline CBF we calculated the average perfusion value across a 30s period prior to the onset of stimulation in the both multifrequency stimulation experimental paradigms (at the start of data collection and after drug administration). A one-way ANOVA on CBF baseline differences (across the saline, scopolamine and donepezil treated animals) found no difference between groups, *F*(2,17)=2.62, *p* =.106. To assess any effects of drug administration upon baseline cortical neuronal activity we computed EEG power over a 30s period of electrophysiological data acquired at the same two time points as the CBF baseline measures. Differences in EEG power were calculated for all five frequency bands and analysed in a one-way multivariate ANOVA. There was no significant main effect of drug group (saline, scopolamine or donepezil) on EEG band power across the 5 bands, *F*(10, 22) = 1.56, *p* =.185; Wilks’ Λ = .343.

### Cerebral blood flow responses to somatosensory stimulation

#### Multifrequency 2s stimulations (1-40Hz)

Repeated measures analysis of variance (ANOVA) was applied to CBF response maxima values in order to determine significant effects of drug administration within each drug group. All cases (*n* = 18) were included in the analyses. All drug conditions had a significant effect of stimulation frequency (saline: *F*(5,25)=7.75, *p*<.001; scopolamine: *F*(5,25)=10.19, *p*<.001; donepezil: *F*(5,25)=21.17, *p*<.001) (Figure 1). Administration of saline did not have a significant effect on the CBF response (*F*(1,5)=5.65, *p*=.063) and there was no significant interaction between drug administration and stimulation frequency (*F*(5,25)=2.07, *p*=.103). Scopolamine or donepezil administration did not itself result in a significant change in the magnitude of CBF responses (scopolamine: *F*(1,5)=0.89, *p*=.390; donepezil: F(1,5)=1.75, *p*=.244), however there was a significant interaction effect of drug administration and stimulation frequency upon CBF response magnitudes for both drugs (scopolamine: *F*(5,25)=4.08, *p*=.008; donepezil: *F*(5,25)=5.15, *p*=.002). Post-hoc analysis of individual stimulation frequencies using paired t-tests indicate significant increases in CBF response magnitudes at 20Hz (*p*=.026) and 40Hz (*p*=.020) for donepezil treated animals (Figure 1B). Scopolamine treated animals showed a marginally non-significant decrease in CBF response magnitude at 40Hz (*p*=.071).

**Figure 1.**
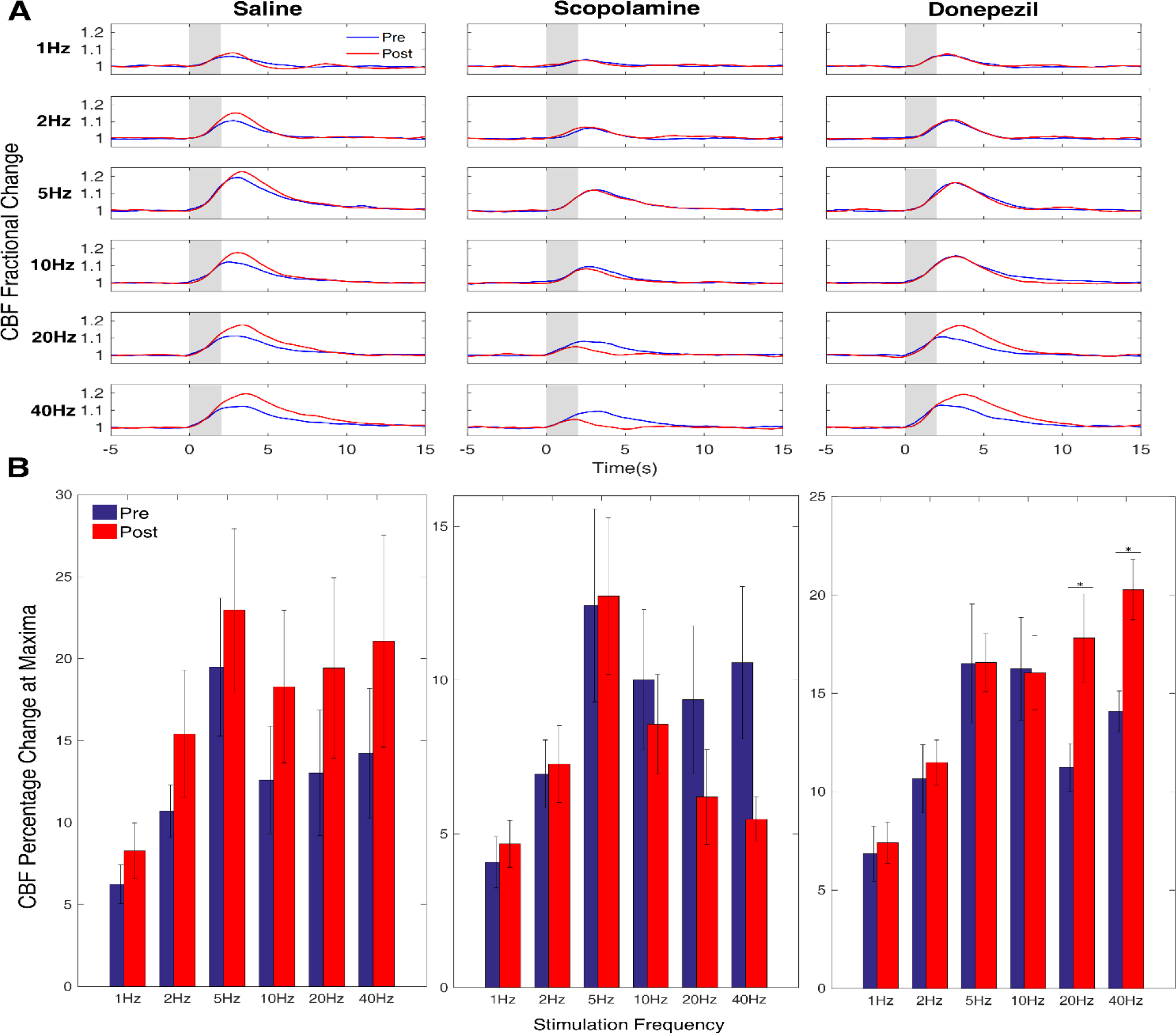
Cerebral blood flow (CBF) responses to 2s whisker stimulation at six frequencies (1-40Hz, 1mA), before (pre, blue) and after (post, red) administration of saline, scopolamine or donepezil. (A) Time series show mean fractional changes in CBF. Grey rectangle indicates stimulation onset/offset and error bars indicate standard error of the mean. (B) Bar charts show change in CBF maxima. *Represents a significant difference at *p*<0.05 between pre- and post-drug treatment for specific stimulation frequencies.

#### 16s long stimulations (10Hz)

Repeated measures t-tests were used to determine significant effects of drug administration on the magnitude of CBF responses to 16s whisker stimulation. An AUC measure was used to quantify responses to 16s stimulation as the maxima is less representative of the longer response profile than that for the short (2s) multifrequency stimulation. Donepezil administration was associated with a significant increase in CBF response magnitude (*t*(5)*= -* 2.64, *p*=.046) (Figure 2 insert), whereas there was no significant effect for saline (*p*=.191) or scopolamine (*p*=.734).

**Figure 2.**
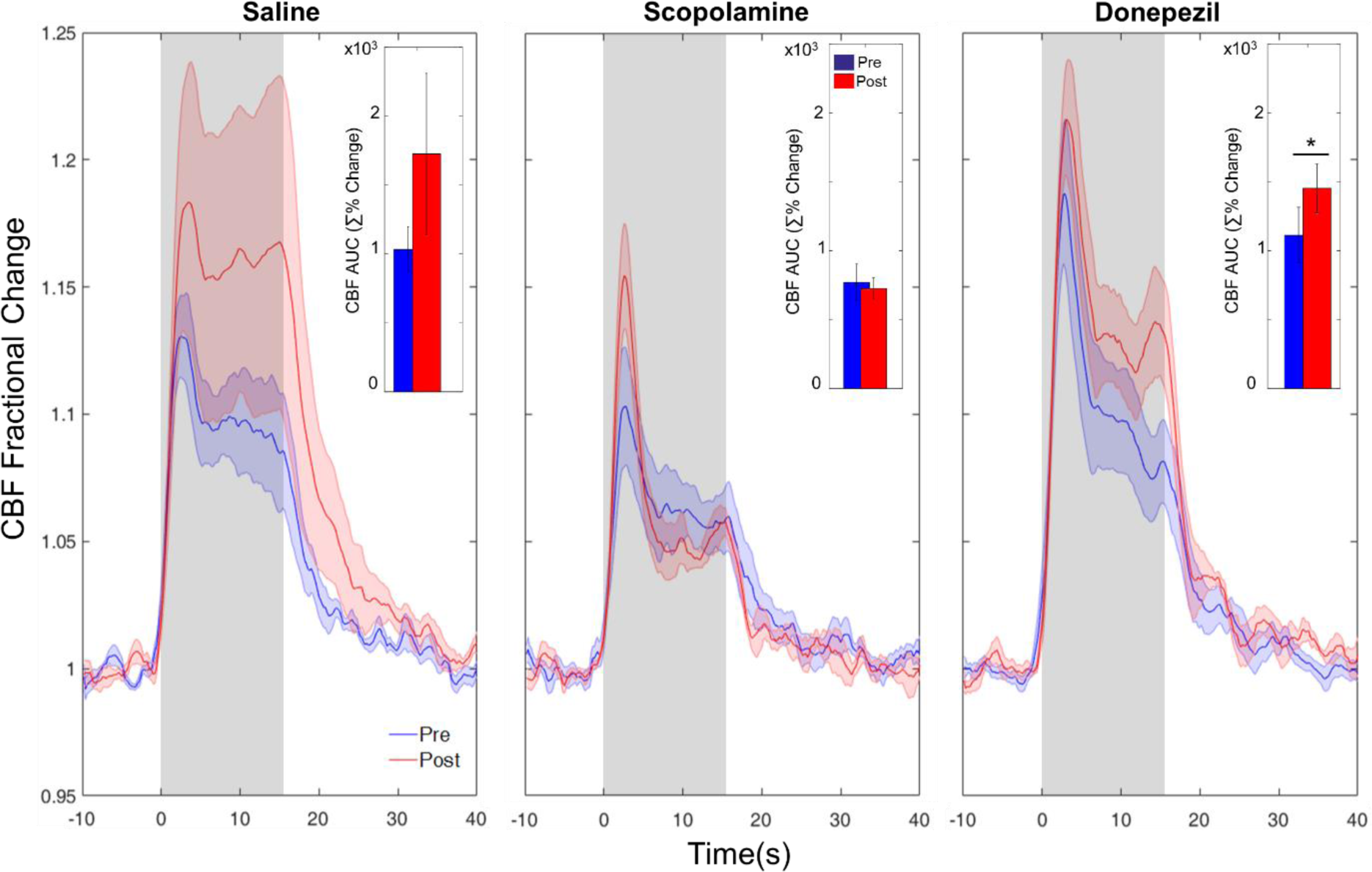
Cerebral blood flow (CBF) responses to 16s whisker stimulation, at 10Hz, before (pre, blue) and after (post, red) administration of saline, scopolamine or donepezil. Time series show mean fractional changes in CBF and bar charts (inset) show change in area under the curve (AUC, units are summed percentage change). Grey rectangle indicates stimulation onset/offset and shaded area represents standard error of the mean. *Represents a significant difference at *p*<0.05 between pre- and post-drug treatment.

### Neuronal responses to somatosensory stimulation

#### Multifrequency 2s stimulations (1-40Hz)

Repeated measures analysis of variance (ANOVA) was applied to the calculated AUC values for the LFP responses in order to determine significant effects of drug administration upon neuronal responses to stimulation within each drug group. All cases (*n* = 18) were included in the analyses. There was a significant effect of stimulation frequency upon LFP response magnitude for all three drugs (saline: *F*(5,25)=11.05, *p*<.001; scopolamine: *F*(5,25)=4.55, *p*=.004; donepezil: *F*(5,25)=29.28, *p*<.001) (Figure 3).

**Figure 3.**
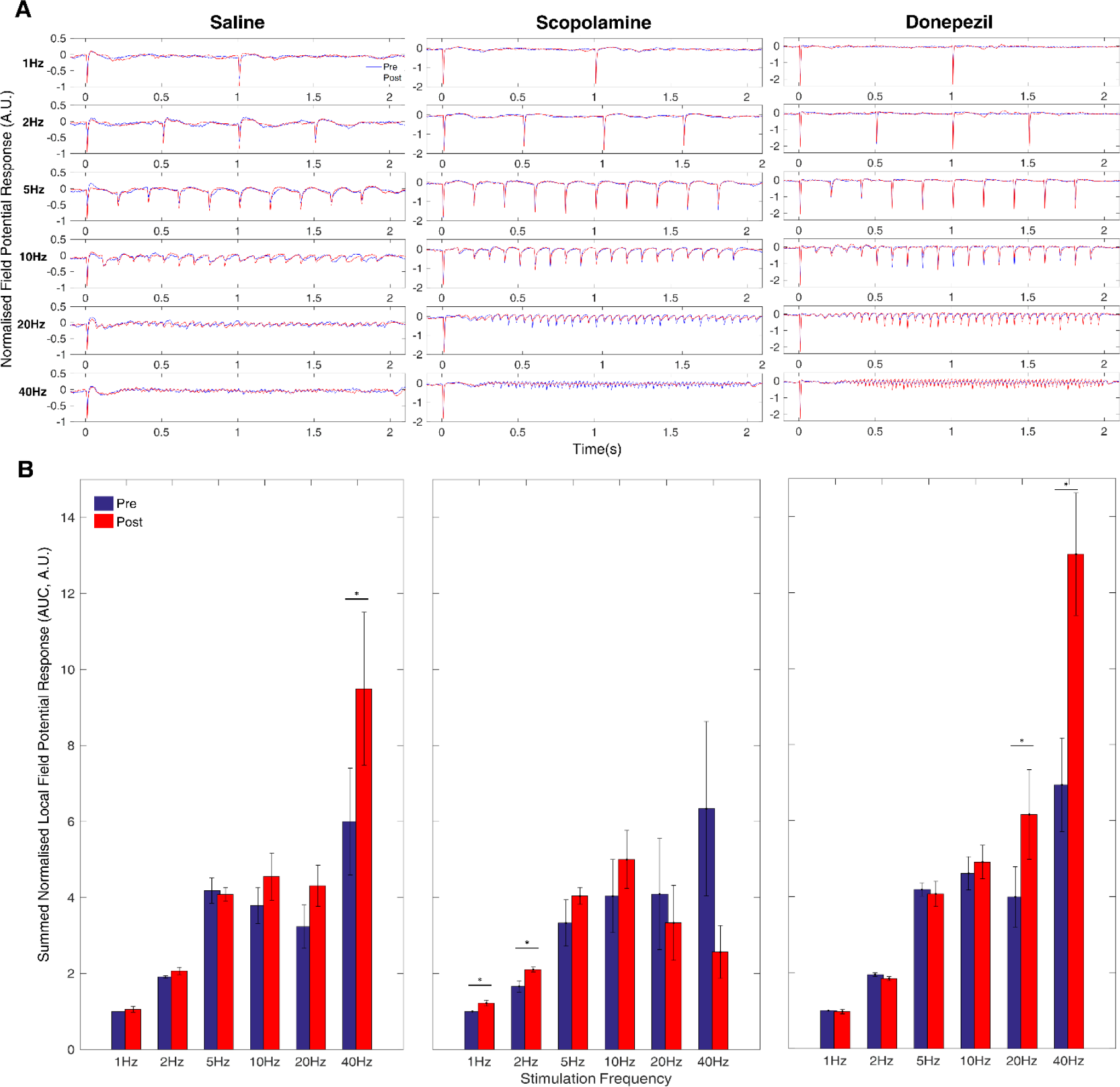
Local field potential (LFP) responses to 2s whisker stimulation at six frequencies, before (pre, blue) and after (post, red) administration of saline, scopolamine or donepezil. Data were normalised to the pre-drug average magnitude of the response to the first stimulus pulse for 1Hz stimulation. (A) Time series show the mean LFP response to the 2s stimulation train. (B) Bar charts show area under the curve (AUC) calculated for each stimulation pulse and then summed across stimulation pulses for each frequency condition. Error bars indicate standard error of the mean. *Represents a significant difference at *p*<0.05 between pre- and post-drug treatment for specific stimulation frequencies.

Administration of saline and donepezil (but not scopolamine) had a significant effect upon LFP response magnitudes (saline: *F*(1,5)=8.85, *p*=.031; donepezil: *F*(1,5)=10.31, *p*=.024; scopolamine: *F*(1,5)=1.34, *p*=.299). There was a significant interaction effect of drug administration and stimulation frequency upon LFP response magnitude for all three drugs (saline: *F*(2,25)=5.9, *p*=.001; scopolamine: *F*(5,25)=5.76, *p*=.001; donepezil: *F*(2,25)=9.32, *p*<.001). Post-hoc analysis of individual stimulation frequencies using paired t-tests indicate significant increase in LFP response magnitude at 20Hz (*p*=.009) and 40Hz (*p*=.026) for donepezil treated animals (as for CBF responses). Scopolamine treatment was associated with an increase in LFP response magnitude at low stimulation frequencies of 1Hz (*p*=.009) and 2Hz (*p*=.007) with a marginally non-significant effect at 40Hz (*p*=.085). Saline treatment produced a marginally significant increase in LFP response magnitude at 40Hz (*p*=.048) (Figure 3B).

#### 16s long stimulations (10Hz)

Repeated measures t-tests were used to determine significant effects of drug administration on the magnitude of LFP responses to 16s whisker stimulation. In line with results for the CBF data, donepezil administration was associated with a significant increase in LFP response magnitude (*t*(5)*= -*5.43, *p*=.003) (Figure 4), whereas there was no significant effect for saline (*p*=.855) or scopolamine (*p*=.112).

**Figure 4.**
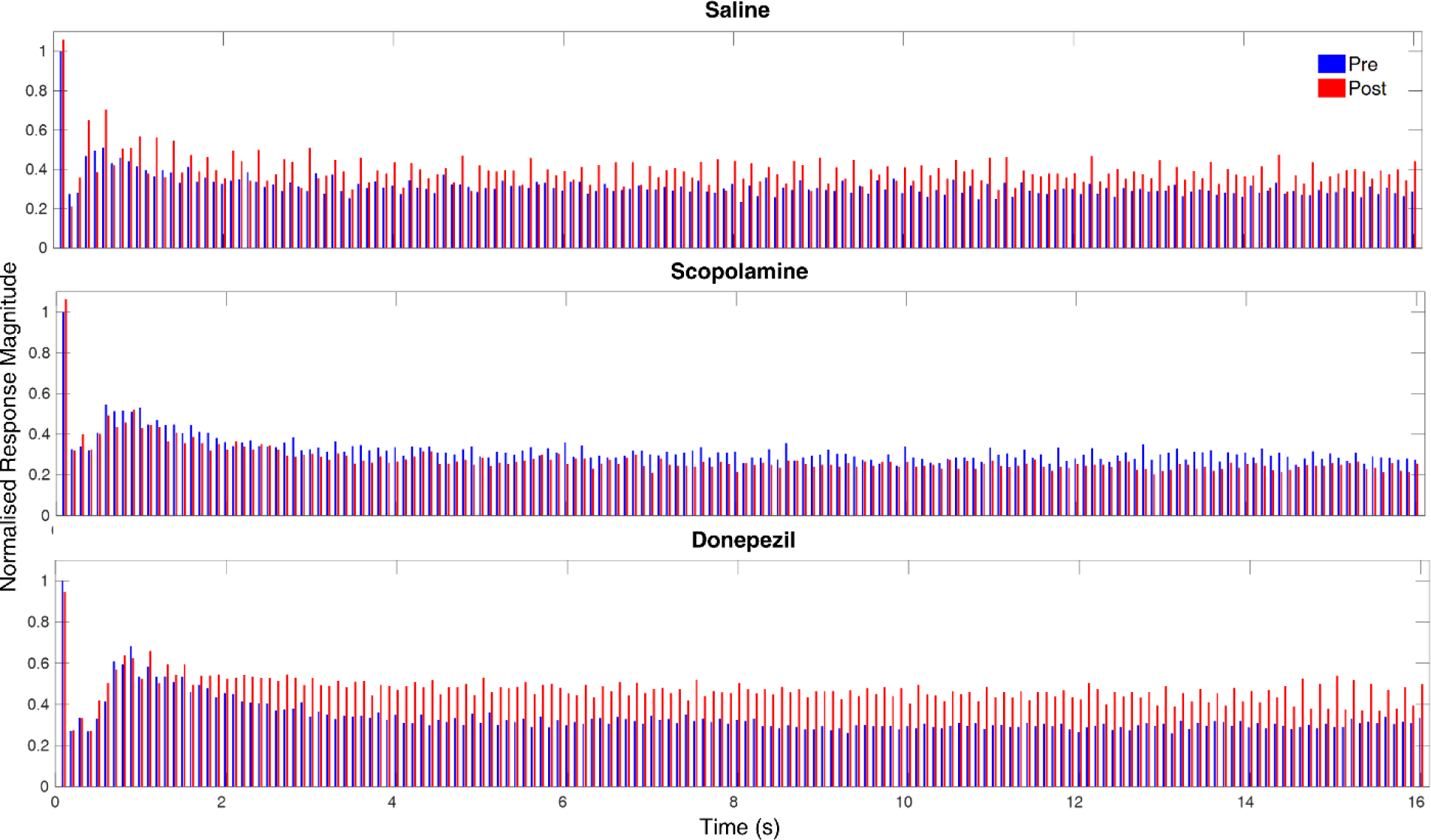
Local field potential (LFP) responses to 16s whisker stimulation at 10Hz, before (pre, blue) and after (post, red) administration of saline, scopolamine or donepezil. Data were normalised to the pre-drug average magnitude of the response to the first stimulus pulse.

#### Neurovascular coupling

To assess neurovascular coupling, data were further analysed on a trial by trial basis by plotting haemodynamic and neuronal response magnitudes against one another (Figure 5). To reduce inter-animal variability each animals’ response values were first divided by their respective mean so as to normalise the datasets. Data from all trials from the multifrequency stimulation experiment were plotted (a total of 720 data points for each drug group). A linear regression was calculated (by least squares) for each data set in order to characterise the neurovascular coupling relationship from the concurrently measured neuronal and haemodynamic responses. The regression line was significantly different from zero in all datasets except in the scopolamine condition after drug administration (Figure 5). T-tests of the difference in the slope of the regression lines revealed a highly significant difference in the scopolamine condition before versus after drug administration (*t*(716) = 3.88, *p*<0.001), indicating an alteration in neurovascular coupling.

**Figure 5.**
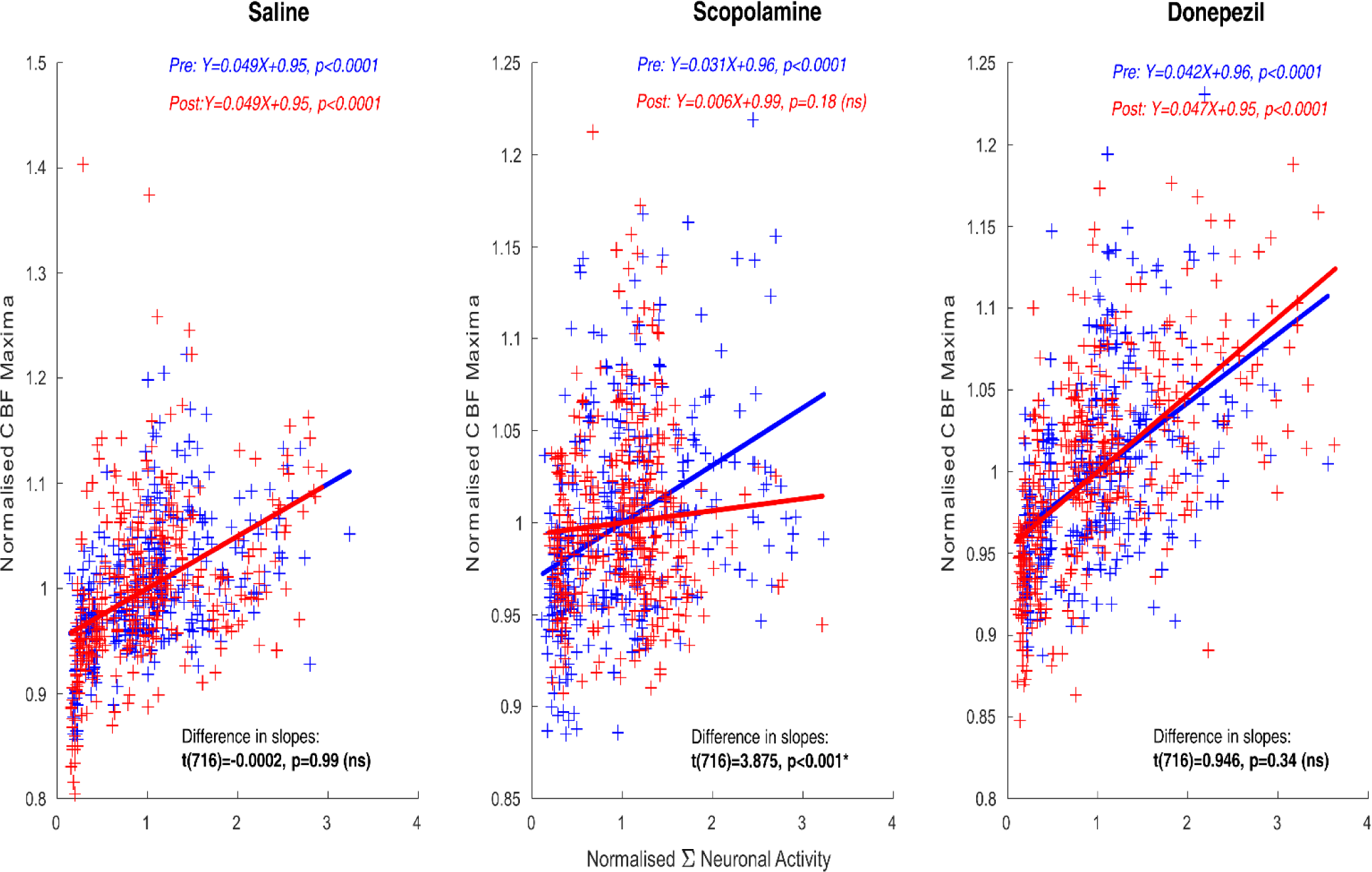
Scatterplots to indicate the relationship between magnitude of neuronal and haemodynamic (CBF) responses. Each data point is from a single trial, for a single animal from the multi-frequency stimulation experiments (60 trials, 10 at each frequency 1, 2, 5, 10, 20 and 40Hz), before (pre, blue) and after (post, red) administration of saline, scopolamine or donepezil. A linear regression model was applied to characterise neurovascular coupling (blue and red lines). The equation of the regression lines are stated as are the results of a t-test of the significant difference of the slope from zero (red & blue text). A further t-test to determine significant differences in the regression models before and after drug administration was conducted (text in black).

## Discussion

The main finding of this study was an alteration of neurovascular responses, including a change in the neurovascular coupling relationship, by pharmacological manipulation of cholinergic function. We have demonstrated haemodynamic and neuronal signal changes in response to scopolamine and donepezil administration. Scopolamine treatment to block cholinergic signalling was associated with an apparent loss of the monotonic neurovascular coupling relationship with no significant dependence of the magnitude of the CBF response on the magnitude of the neuronal response. These findings have implications for neuroimaging data acquired from clinical cohorts that have reduced cholinergic function such as AD patients (*30*), as well as for the use of neuroimaging to study brain effects of cholinergic manipulations of cognitive function (*31*). Caution should be exercised when interpreting human fMRI data from these groups as these data rely on an assumed monotonic neurovascular coupling function (*24*) where haemodynamic and neuronal response magnitude change in tandem with one another. Our findings furthermore highlight that inferences about drug effects on neuronal activity cannot necessarily be extrapolated from haemodynamic neuroimaging signals. Moreover, there is a need for improved understanding of neurovascular consequences of pharmacological manipulations of neurotransmitter systems, which may require further studies that concurrently measure both haemodynamics and neuronal activity.

A significant interaction effect of stimulation frequency and either donepezil or scopolamine administration upon CBF response magnitudes suggests an altered relationship between the magnitude of stimulus input (manipulated here via stimulation frequency) and the magnitude of the associated haemodynamic response. An altered response pattern is qualitatively indicated by inspecting the bar charts plotting response magnitudes at each stimulation frequency (Figure 1). Similar effects are apparent for the neuronal response data (Figure 3). Comparing across the neuronal and haemodynamic data sets is challenging, a problem avoided in studies which focus on a single stimulation protocol in an attempt to determine whether neurovascular relationships are altered in a particular context. Our data highlight the underlying complexity of neurovascular coupling relationships, where the effects of pharmacological or disease related manipulations could be dependent on the properties of the stimulus protocol employed.

By plotting trial by trial pre and post cholinergic challenge haemodynamic and neuronal data together to assess neurovascular coupling more directly, a clear pattern of neurovascular alteration or uncoupling was evident for the cholinergic antagonist drug scopolamine. This was not apparent when analysing haemodynamic and neuronal activity data separately due to the effect of trial averaging in the methods to visualise these data. The difference in the linear regression model before and after administration of scopolamine demonstrates an alteration of neurovascular coupling following pharmacological antagonism of brain acetylcholine by scopolamine, where stimulus evoked changes (increases or decreases) in neuronal demand are not accompanied by concomitant changes in CBF.

An equally important finding from this study was that the widely prescribed cholinergic agonist donepezil does not, at the dosage used here, affect the relationship between concurrently measured neuronal activity and CBF changes. This suggests that changes in the magnitude of haemodynamic (including fMRI) signals in human or preclinical studies in response to cholinergic agonists do reflect changes in underlying neuronal activity, rather than effects on, for example, vascular reactivity. Coupled with the finding of Kocsis et al. (2014) which suggests the effects of cholinergic agonists on cognitive function are due to vascular drug actions (as drug BBB penetration was shown not to be necessary), this raises the possibility that modulation of cortical vascular function can impact directly on neuronal function (the hemo-neural hypothesis, (*32*). However, a limitation to the present work is that it was carried out in healthy animals and it will be important in future research to determine if donepezil treatment is protective of neurovascular function under conditions of cholinergic loss/blockade and furthermore whether such effects require BBB drug penetration.

Our findings are in good agreement with Lecrux et al. (2017) who also recently examined cholinergic modulation of neuronal and haemodynamic responses and reported altered neurovascular function as well as augmented neuronal and haemodynamic responses following treatment with cholinergic agonists. We do not find a pronounced reduction in the magnitude of haemodynamic responses following scopolamine administration (some reduction is indicated for higher stimulation frequencies), although comparison of the dosing for scopolamine between these studies is complicated because of the differing routes of administration (intravenous vs. intracisternal). An important difference between the studies is the use of shorter stimuli with variation in stimulus intensity (frequency) in the present work so as to reveal any changes in neurovascular coupling over a wider dynamic range of neuronal inputs. In this context we find that the magnitude of the haemodynamic responses no longer provides information regarding the magnitude of the underlying neuronal response because it is not clear from the trial-averaged measures of neuronal and haemodynamic response magnitude (Figures 1 & 3) that neurovascular coupling is substantially altered after scopolamine administration. We suggest that potentially important effects of experimental manipulations upon relationships *between* neurophysiological parameters may be masked by trial averaging, where such features are discarded as noise.

We observe increases in both CBF and neuronal response magnitudes following treatment with donepezil, with this finding evident in both the short (2s) and long duration (16s) experimental conditions. It has been hypothesised that similar observations of increased CBF observed with donepezil treatment in human studies may be related to increases in neuronal activity (*33*), maintenance of adequate CBF supply (*34, 35*) or normalising of flow regulation. As a result, improved blood supply to active neurons in the AD brain (*36*), may underlie some of the restoration of cognitive performance afforded by such pharmacological interventions (*37*). Although we do not find an effect of drug treatment upon baseline CBF, our neurovascular measurements appear to support this hypothesis with evidence that acute pharmacological *decreases* in brain acetylcholine may diminish the capacity of CBF to match changing stimulus-evoked neuronal demands. However, we do not find evidence that (acute) cholinergic agonist treatment leads to an equivalent ‘oversupply’ of CBF with respect to neuronal demands: we hypothesise that such an effect will emerge if the prerequisite cholinergic tone for normal vascular responses (*14*) is not present, such as under disease conditions.

We are unsure as to why CBF and neuronal response effects of scopolamine or donepezil treatment appear to be manifested at differing frequencies of sensory stimulation, although similar effects have been recently observed for serotonergic manipulation (*24*). In addition, comparisons of responses in awake and anaesthetised animals (*38*) to whisker stimulation reported response magnitude differences between awake and anaesthetised animals that were stimulation-frequency dependent, with more pronounced differences in response magnitudes evident at higher stimulation frequencies. Since acetylcholine is a potent neuromodulator playing an important role in arousal and attention, it is possible that the cholinergic challenge used here led to a potentiation of the cortical response to higher sensory stimulation frequencies, as found in awake animals (*39*), although we do not find evidence for an alteration in baseline cortical neuronal activity.

In conclusion we provide evidence that modulation of brain acetylcholine alters the relationship between neuronal and haemodynamic responses, with a neurovascular uncoupling effect associated with reduced cholinergic function. This finding has implications for the interpretation of BOLD fMRI and BOLD phMRI data in both animal and human imaging studies where cholinergic function may be different between experimental or patient groups. Our findings support the notion that reduced cholinergic modulation of neurovascular function may be an important component of neurodegenerative diseases such as AD highlighting a role for cholinergic drugs in normalising neurovascular function.

## Acknowledgements

The authors would like to thank Dr Aisling Spain for assistance with the experimental work and development of the experimental protocol.

## Authors’ Contribution Statement

CM and GB designed the experiments, GB performed the experiments, GB and CM analysed the data, GB, CM and JB interpreted the data, GB, CM and JB wrote the manuscript.

